# Tutorial on quantifying and sampling biomolecular ensembles with ShapeGMM

**DOI:** 10.1101/2025.10.06.680711

**Authors:** Subarna Sasmal, Martin McCullagh, Glen M. Hocky

**Author notes:** Electronic mail.

## Abstract

Here we present a detailed workflow for clustering and enhanced sampling of biomolecular conformations using the ShapeGMM methodology. This approach fits a probabilistic model of biomolecular conformations rooted in the idea that the free energy can expressed in terms of local fluctuations in atomic positions around metastable states. We demonstrate using a single model system how to generate and fit equilibrium molecular dynamics simulation data. We then show how to use the resulting model to generate a reaction coordinate between two states, how to sample along that coordinate using Metadynamics using our size-and-shape PLUMED module, and how to cluster those biased conformations to give a refined equilibrium ShapeGMM model.

## I. INTRODUCTION

It is now commonplace to sample hundreds of thousands or millions of molecular snapshots by performing molecular dynamics (MD) simulations of biomolecules.^1^ Given the large amount of data, clustering is a valuable, and often overlooked step in performing an unbiased analysis of the output.^2,3^ There are a plethora of clustering algorithms and features to choose from and they all have their pros and cons as well as intricacies to implementation.

ShapeGMM, developed three years ago, is an appealing approach because it marries a physically motivated feature choice—atomic cartesian positions—with fitting a probabilistic/generative model for the data—a Gaussian Mixture Model.^4^ As described below, this works by considering the system in so-called “size-and-shape” space, where configurations invariant to rotation and translation are considered equivalent, which allows the use of particle positions as features.^4^ The use of a probabilistic model provides a number of additional benefits to heuristic clustering, such as *k*-means, as it allows for the estimate of Thermodynamic properties, such as configurational entropy, to quantify the ensemble.^5^ Moreover, we have demonstrated that *equilibrium unbiased* ShapeGMM models can be computed from *biased* sampling methods such as Metadynamics (MetaD),^6^ meaning that it is possible to characterize a conformational ensemble which can only be effectively sampled using enhanced sampling approaches.^5^ Finally, we demonstrated that reaction coordinates computed to separate different states in the ShapeGMM model can be effective for sampling between those states, and that these coordinates can be improved through iteratively training on biased sampling along them.^7,8^

In this tutorial, we briefly recap the theory behind fitting a ShapeGMM model, including discussion of good practices for picking the number of clusters.^4^ We also describe the computing of Linear Discriminant Analysis (LDA) coordinates in size-and-shape space,^7^ and how to sample along these coordinates with well-tempered metadynamics (WT-MetaD).^6^

Although we have applied ShapeGMM on much more complex systems including the cytoskeletal protein actin and a viral helicase,^5,9,10^ here we demonstrate all of the procedures we have developed on the prototypical alanine dipeptide (ADP) in vacuum. For ADP, it is possible to generate all of the data used in this tutorial and perform all of the analysis within a matter of a few hours, something that would not be possible for more complex examples. Sufficient detail is given that adapting our procedure to another system requires only minor modifications, although one will need to consider computational cost depending on system size (see Appendix VII). We also provide, in an accompanying software repository, GROMACS^11^ and PLUMED^12^ inputs needed to generate the data; all the generated data needed to run the analysis steps of this procedure; worked-out python Jupyter notebooks for all parts of the tutorial; and, Docker/Singularity containers with pre-installed software that allow all steps to be exactly reproduced (see Sec. VI).

The rest of the tutorial is organized as follows: in Sec. II we give the simple procedure for installing python and PLUMED modules for performing ShapeGMM analysis and enhanced sampling using coordinates defined in size-and-shape space; Sec. III gives the theoretical background for the rest of the article; Sec. IV contains our step-by-step tutorial; finally, in Sec. V we conclude with final thoughts on our implementation and its uses.

## II. INSTALLATION

Python code to perform ShapeGMM, as well as a helper module for performing weighted linear discriminant analysis are available on GitHub.^13,14^ To install these modules, one can simply run:

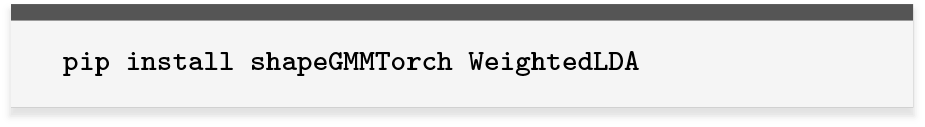

`ShapeGMMTorch` makes use of the `PyTorch` library in python,^15^ and will be automatically installed. To ensure that your setup can make use of GPU acceleration on an available NVIDIA graphics card, one should first install `PyTorch` with CUDA following their instructions.

Finally, to install PLUMED with our size-and-shape module, one can simply run:

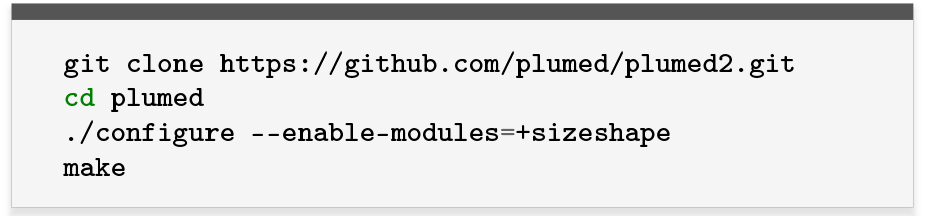

GROMACS can be patched with PLUMED and installed using the instructions on the respective websites, which are beyond the scope of this tutorial.

Installed versions of these software suitable for performing the steps of this tutorial and other research projects are provided in Docker/Singularity containers, as described in Sec. VI.

## III. THEORY AND METHODS

ShapeGMM is a Gaussian Mixture Model (GMM) clustering algorithm specifically designed for particle positions. The ShapeGMM probability density is given by

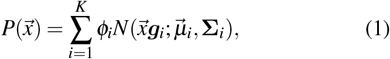

where *K* is the number of components, *ϕ*_*i*_ is the weight of each component with normalization restriction 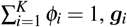 is the rotation matrix that minimizes the Mahalanobis distance between 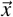 and 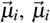 is the *N* × 3 array of the mean position of component *i*, and **Σ**_*i*_ is the 3*N* × 3*N* covariance matrix for component *i*. The quantity 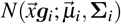 is the value of the multivariate normalized Gaussian of component *i* evaluated at position 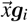, and is given by

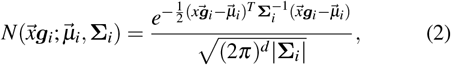

where *d* is the rank of **Σ**.

ShapeGMM employs an expectation maximization (EM) algorithm coupled with maximum likelihood structural alignments to maximize the log-likelihood of the model as a function of the parameters (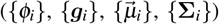) given a data set. The log-likelihood of a data-set of *M* frames is

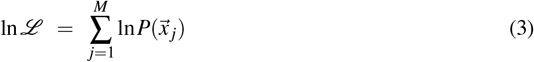

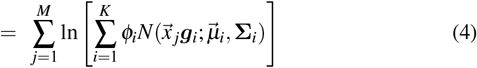

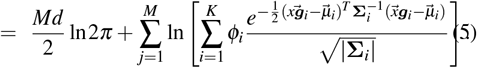

Practically, we do not include the 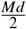 ln 2*π* term in the code because it is independent of parameters and we also report the log-likelihood per frame (*M*^−1^ ln ℒ) so that it is reasonable to compare fits with different training set sizes.

### A. Choosing the covariance model

For most simulation representations of biomolecules, one would expect the covariance matrix of each component to be unrestricted other than the requirements that it be symmetric positive semidefinite of rank 3*N* − 6. Due to the need for structural alignments in ShapeGMM, however, we must restrict the covariance model further; alignment algorithms are only known for a restricted subset of covariance matrices. It is not, however, atypical to consider restricted covariance models in other applications of GMMs.^16^

The most common alignment procedure, based on the Kabsch algorithm,^17^ is to equally weight each atom, particle, or alignment landmark. This is equivalent to considering the covariance to be proportional to the identity matrix,

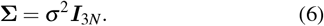

We call this covariance model `uniform`. Assuming this covariance model, the mean and variance of a data set can be readily determined in a maximum likelihood procedure. We have found, however, that this covariance model struggles to differentiate components in molecular models with regions of heterogeneous variance. This could be a protein with a comparable sized globular and unstructured regions, for example. In such cases we recommend using a less restrictive covariance model in the code we call the `kronecker` model in which

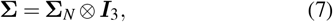

where **Σ**_*N*_ is the *N* × *N* covariance and ⊗ is the Kronecker product. A maximum likelihood alignment procedure using this covariance model will iteratively determine the mean and *N* × *N* covariance of a data set.^4^

There are additional considerations when choosing between the `uniform` and `kronecker` covariance models. First, a `uniform` alignment is much less computationally demanding than a `kronecker` alignment. Second, and related, the number of parameters being fit in the `kronecker` model is much larger than that of the `uniform` model. The `kronecker` covariance has 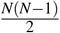 parameters while the `uniform` variance is a single parameter. For these two reasons, it may be impractical to converge the `kronecker` covariance model for large systems (i.e. using many hundreds of atoms, although one can use a coarse-grained representation).

It should be noted that because the likelihood depends on the inverses of the covariance matrices, also called the precision matrix, shapeGMM stores only these matrices. The covariance matrices can be reconstructed by computing the pseudo inverse of the precisions.

### B. Picking the number of clusters

The only hyperparameter of the model is the number of components, *K*. If this number is not known *a priori*, we suggest scanning the number of components and using a combination of the elbow heuristic and cross-validation. For the elbow heuristic, we look at the log-likelihood as a function of number of components in the training data. As the number of components increases, and considering the same training set, the log-likelihood should monotonically increase. The elbow in this plot represents the number of components above which increasing the number of components does little to improve the fit.

Cross-validating the fit allows us to assess whether the model is truly predictive or overfit. This is achieved by dividing a data set into a training portion and a cross-validation portion. The model is then fit on the training portion and the log-likelihood per frame is assessed on both portions. If the log-likelihood per frame is drastically different between the training set and the cross-validation set this indicates over or under-fitting. In a component scan, you want to choose a number of components for which the model is not over or under fit.

`shapeGMMTorch` has a built-in utility to perform this scan (shown below) and a built-in plotting function to plot the results (see accompanying Jupyter notebooks).

### C. Thermodynamic quantities that can be computed from probabilistic models for a configurational ensemble

A ShapeGMM probability density can be used to compute different equilibrium thermodynamic quantities for conformational ensembles of a biomolecular system. Here, we discuss about few such quantities below,

#### 1. Configurational Entropy

The estimation of configurational entropy of solute molecules has been of significant interest over the years.^18–24^ Recent methods have focused on using either the quasiharmonic approximation on particle positions^25,26^ or mutual information on internal coordinates^22–24,27^ to estimate configurational entropy. Here, we estimate the entropy of the shapeGMM mixture of multivariate Gaussians using Monte Carlo integration to approximate the integral,

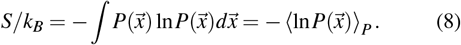

ShapeGMM has an in-built `generate` function that produces configurations sampled from the underlying probability distribution that is obtained from the ShapeGMM fit of training data. The samples are generated in accordance with the relative probability of different metastable states, as computed by ShapeGMM. We note that this is not the true absolute configurational entropy so only entropy differences should be reported.^24^

#### 2. Nonequilibrium Relative Free Energies

Another useful quantity is the relative entropy or Kullback-Leibler divergence that estimates the work to move from one distribution *Q* to another distribution *P*. It is given by

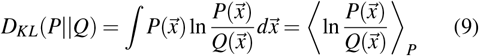

Similar to the case of measuring configurational entropy, here again, the configurations are sampled from distribution *P* using the `generate` function and the average value of the differences in log-likelihoods of these newly generated samples for distributions *P* and *Q* is computed. KL divergence is a non-equilibrium free energy estimate and is not necessarily symmetric.^28^ For comparing distance between clusters, we instead implemented and use the symmetric Bhattacharya distance, as shown in Sec. IV F.

### D. Metadynamics

MetaD works by adding a bias constructed through periodic addition of Gaussian functions centered at the current position of the system in CV space.^6^ In WT-MetaD, the bias height decreases exponentially with the amount of bias already applied at the current value of the CV, *s*(*t*). As a result, the bias *V* is given by the expression

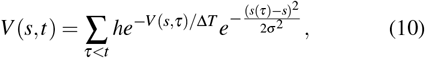

where *h* is the initial hill height, *σ* sets the width of the Gaussians, and Δ*T* is an effective sampling temperature for the CVs. In PLUMED, rather than setting Δ*T*, we choose the bias factor *γ* = (*T* +Δ*T*)/*T*.^6^ In this work, we then reconstruct the free energy surface (FES) and fit models from weighted data by assigning each snapshot a weight.

### E. Using bias weights

In our recent work, we have demonstrated that ShapeGMM can be used to obtain the unbiased estimates for conformational ensembles of a biomolecular system by training directly on the enhanced sampling trajectories obtained from Metadynamics simulations. A detailed description on choosing the correct choice of weights and computing them for training samples from the biased data are provided in that work.^5^ For the purpose of this tutorial, we are going to use weights which are given by,

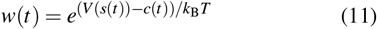

where 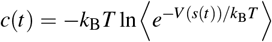 is an average accumulated bias over the CV grid at a fixed time.^6,29^ This so-called “reweighting bias” or ‘rbias’ can be computed and reported automatically in PLUMED.^30^

## IV. TUTORIAL

### A. Generating the input data

Here, we generate 20 ns of data starting from each of two configurations of ADP in vacuum, with inputs adapted from the PLUMED-tutorials resource.^31^ We save data every picosecond, resulting in 20K configurations for each state.

Due to changes in how newer versions of GROMACS handle vacuum simulations, to generate the data as done in this example, one must use a version of GROMACS-2019 patched with a modified version of PLUMED 2.8.4. This is available in the provided singularity/docker container (Sec. VI). We note that a second singularity/docker image is also provided that includes GROMACS 2024.3 with PLUMED 2.10b, appropriate for other examples.

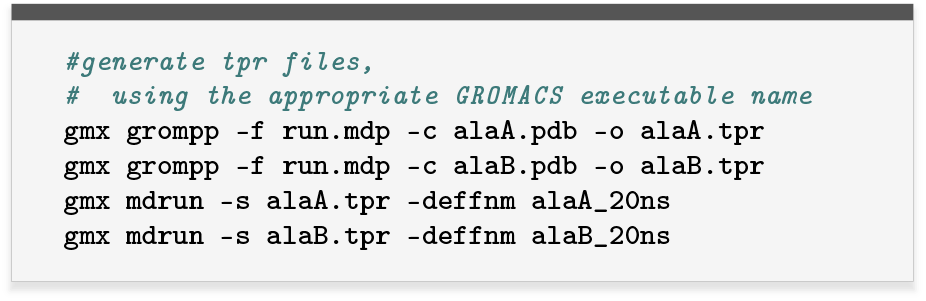

We can also generate a reference well-sampled conformational landscape by biasing the *ϕ* and *ψ* backbone dihedral angles using WT-MetaD as described in the PLUMED tutorials^31^ (Fig. 1), as shown in the abbreviated example below.

**FIG. 1.**
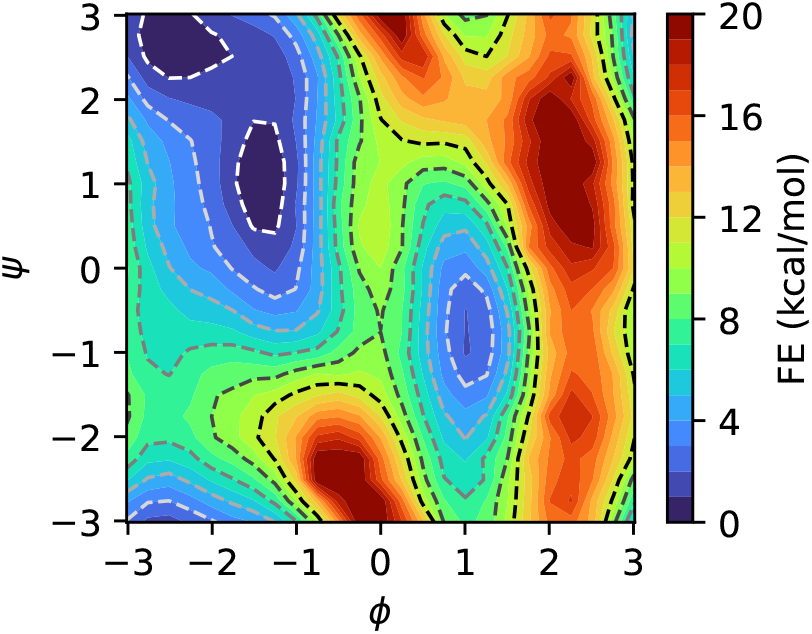
Reference free energy surface for ADP computed by WT-MetaD on backbone dihedrals.

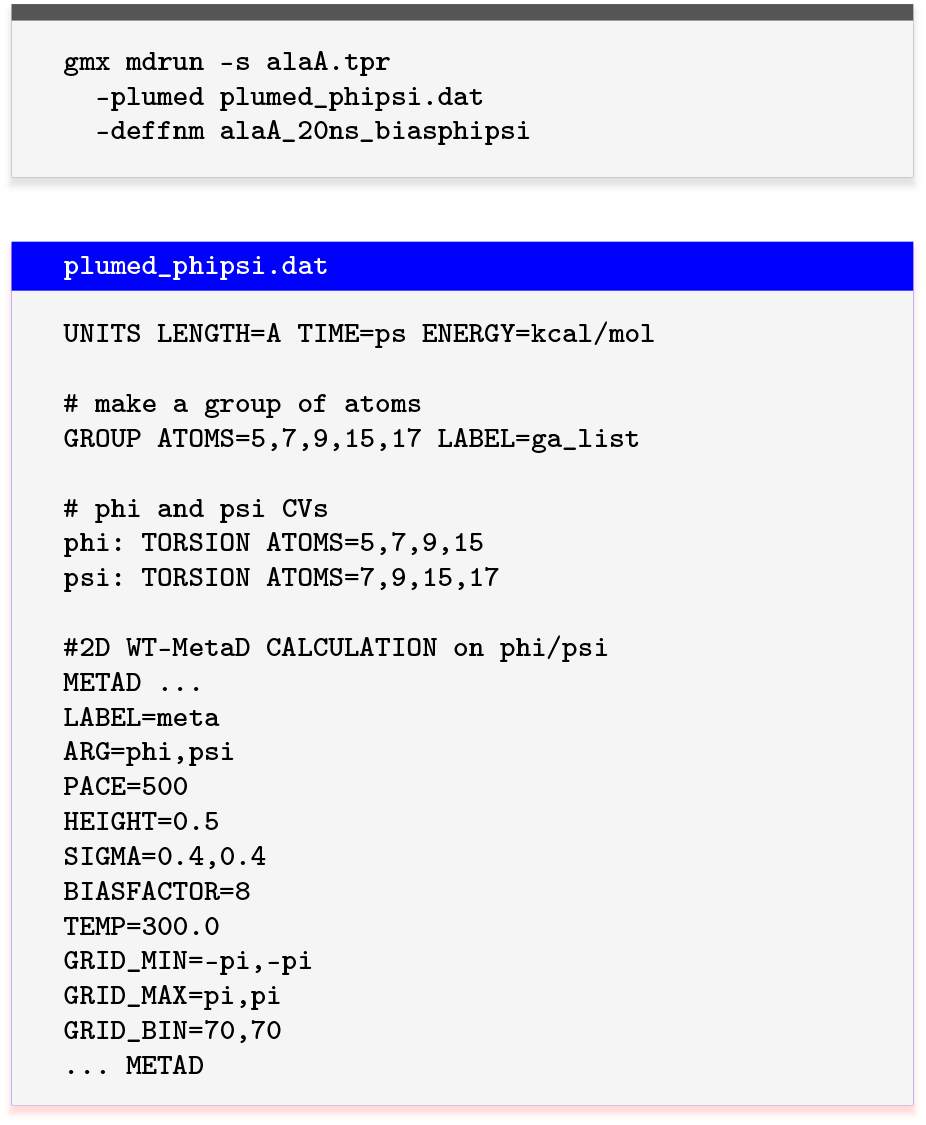

### B. Reading and formatting the data for shapeGMMTorch

To perform clustering, we need to first read and process MD trajectory data, and select the atoms we want to use as input features. Here, we give an example of how to perform this for ADP in our two states using MDAnalysis.^32^ The result is a numpy^33^ array of dimension *M* × *N* × 3, where *M* is the number of frames and *N* is the number of chosen atoms/particles.

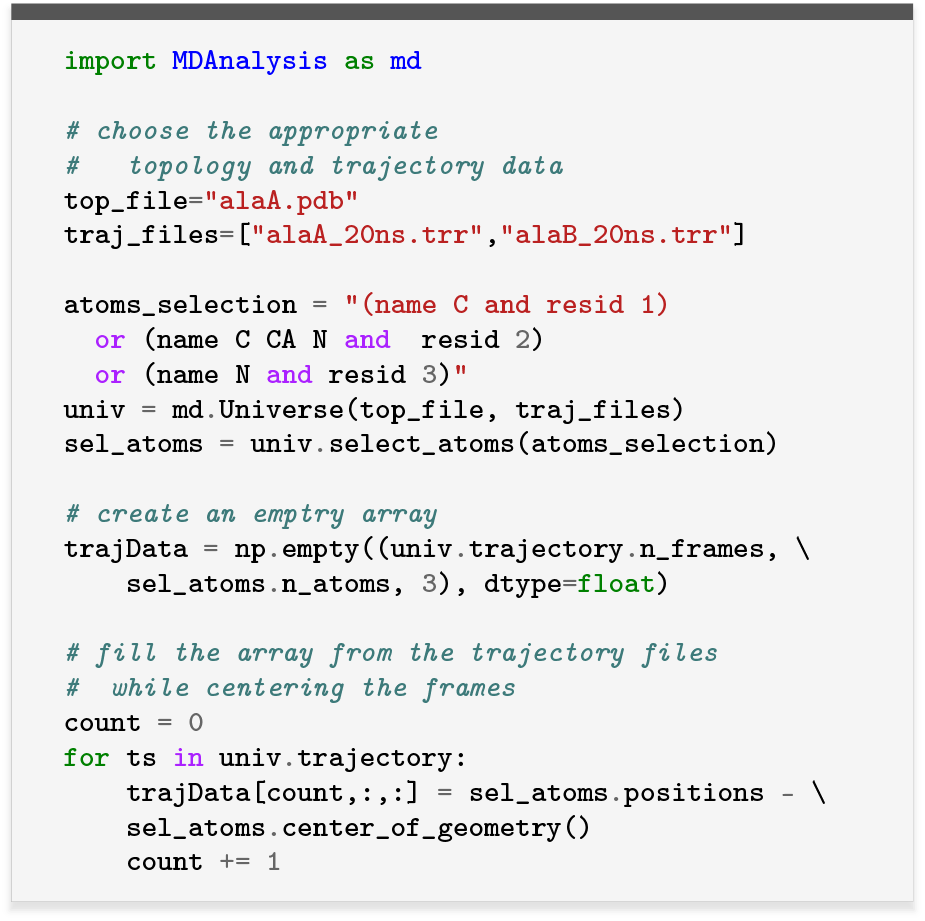

In this case, after reading in the data, we can compute the backbone dihedral angles for the peptide and show that configurations from our two simulations remain in their starting states (Fig. 2).

**FIG. 2.**
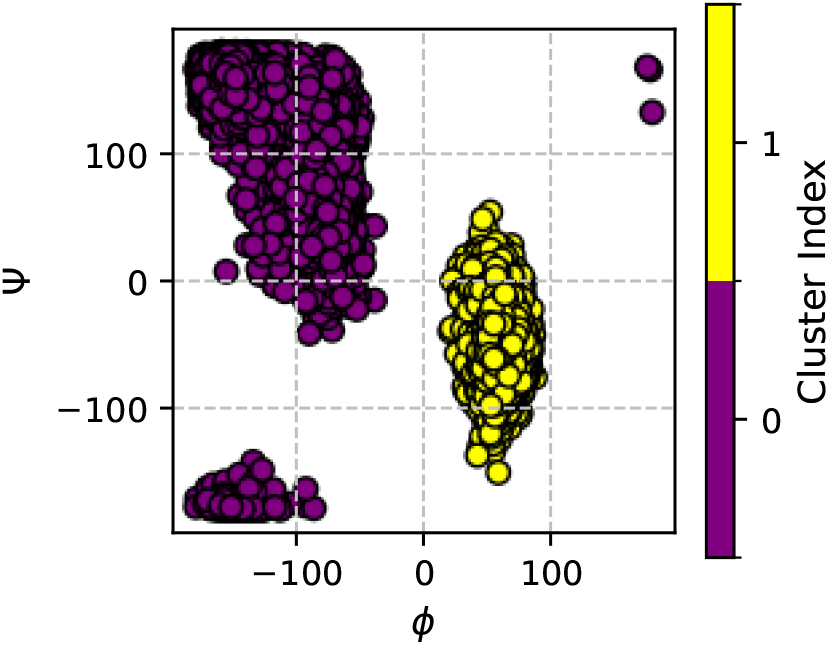
Ramachandran plot showing data from two 20ns simulations of ADP in vacuum, colored by starting configuration.

### C. Performing clustering with shapeGMMTorch

We have consolidated the most typical protocol for clustering, including scanning the number of clusters *K* and performing cross validation and log-likelihood calculations into a single function. The result is shown in Fig. 3.

**FIG. 3.**
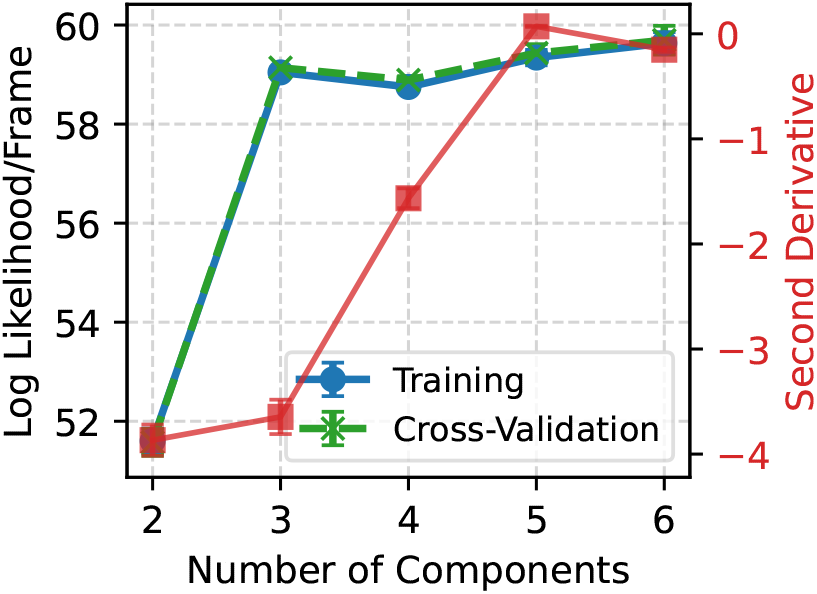
The maximized log likelihood is shown for each number of clusters *K*, with blue showing the result for training data and green for validation set. The overlap indicates a lack of over-fitting. In red, we show the second derivative of the training log-likelihood; the large jump in likelihood combined with a minimum at *K* = 3 suggests choosing 3 clusters.

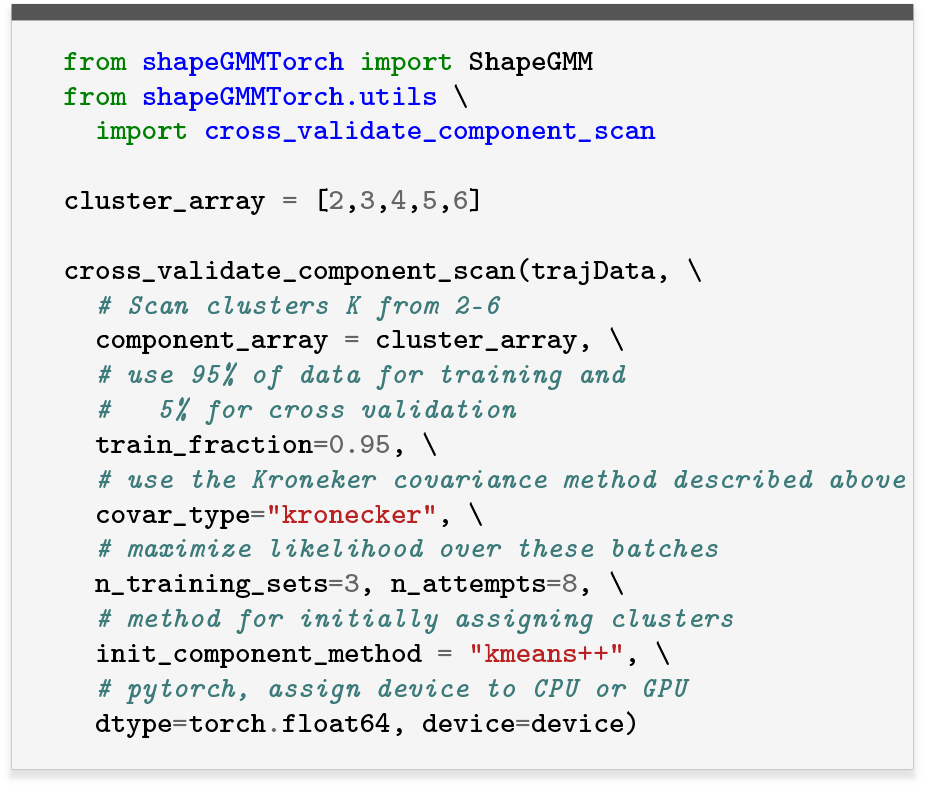

Results from this scan indicate *K* = 3 is the best choice of clusters, so we next proceed to cluster all the data using a ShapeGMM model with *K* = 3 components. As can be seen in Fig. 4, the approach properly identifies two sub-states within the upper left basin, as expected from the reference free energy surface.

**FIG. 4.**
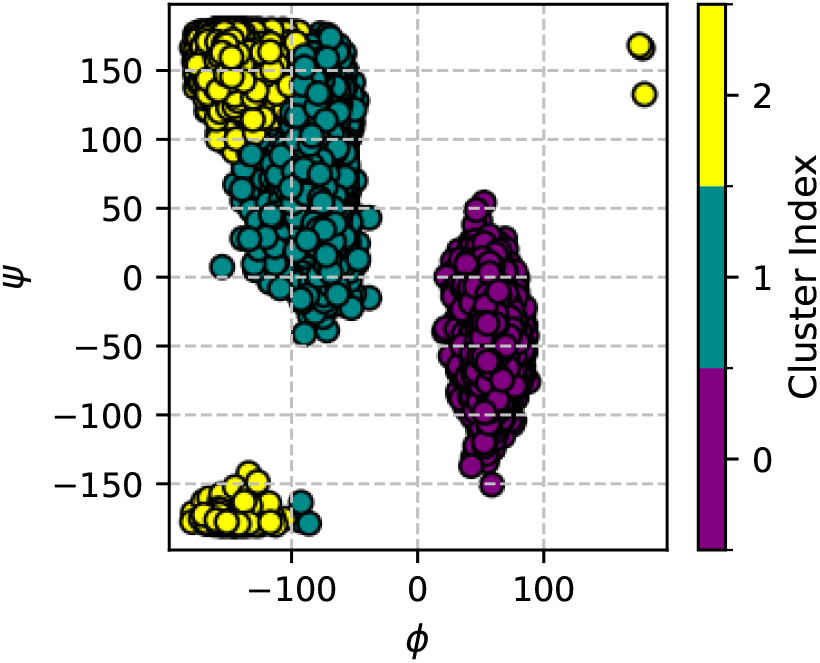
Ramachandran plot with points colored by *K* = 3 clustering on the data shown in Fig. 2.

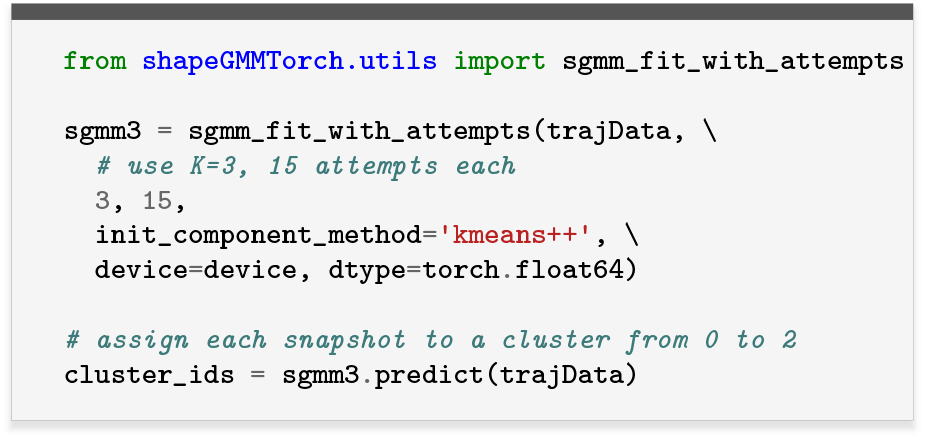

### D. Generating a reaction coordinate with Position LDA

To make a reaction coordinate between cluster 0 (purple) and 1 (green) in Fig. 4, we first iteratively align the frames in the two separate trajectories to a common global mean/covariance. We then extract the frames belonging to cluster 0 and 1.

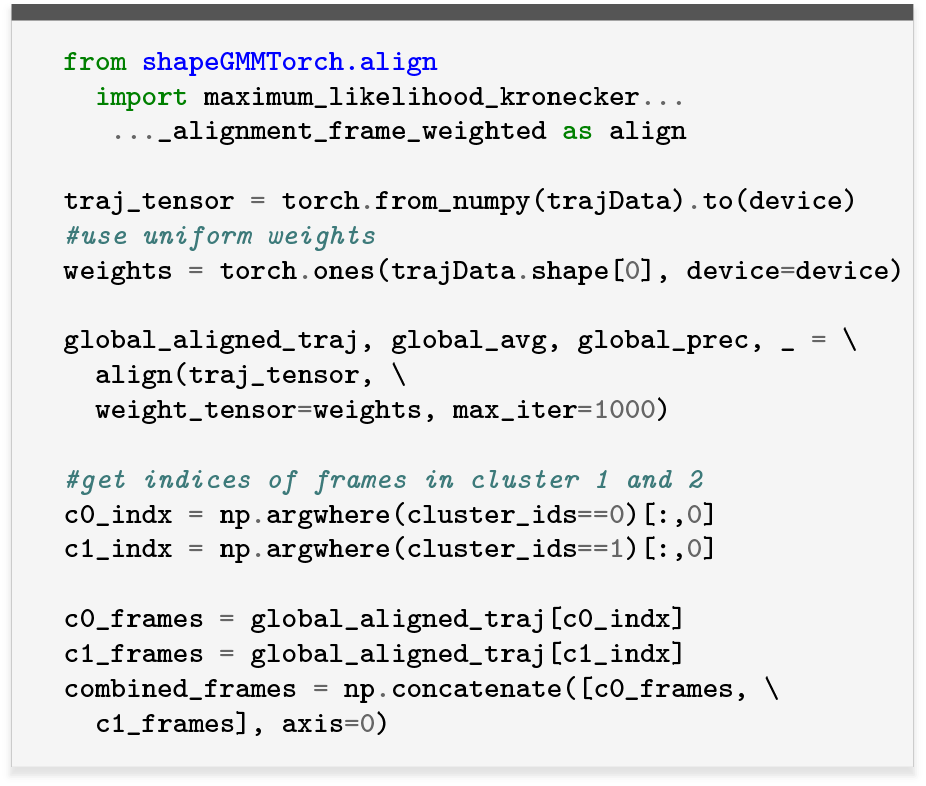

Finally, we use the WeightedLDA library to compute an LDA coordinate that creates a linear projection of the input atomic coordinates to best separate these two clusters. The syntax is based on the LDA method in scikit-learn.^34^ The output values are shown in Fig. 5

**FIG. 5.**
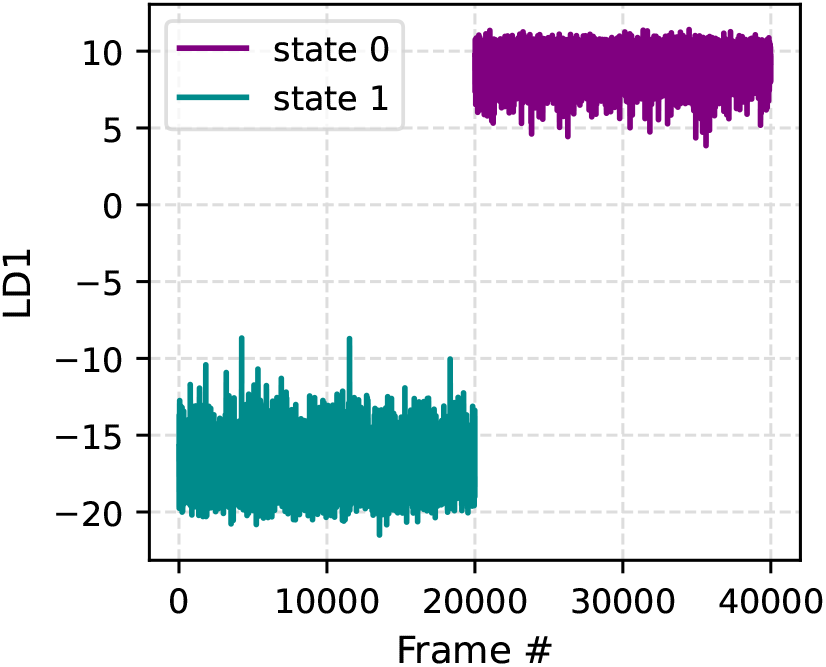
LD value for each frame after performing fitting on globally aligned data, for clusters 0 and 1 in Fig. 4.

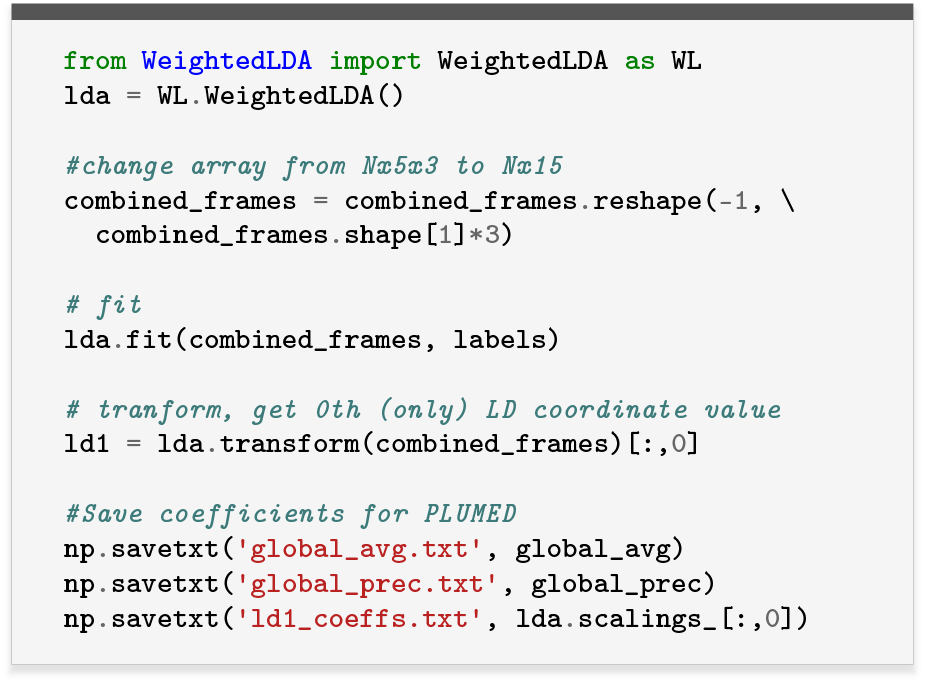

### E. Enhanced sampling with the PLUMED SizeShape module

The `sizeshape` PLUMED module adds two CVs to PLUMED: our LDA CV (`SIZESHAPE_POSITION_LINEAR_PROJ`) and the Mahalanobis distance of the current configuration to the center of a Gaussian considering a particular covariance (`SIZESHAPE_POSITION_MAHA_DIST`). Only the first of these is used in this tutorial. We note that in our original implementation, `SIZESHAPE_POSITION_LINEAR_PROJ` was named `LDA_PROJ` and the argument `COEFFS` was termed `VECTOR`.^7^ The older form is used in the provided input files for this tutorial, but the newer one is shown here in the main text and should be used for any future studies.

Using coefficients from the previous section, we run new MetaD simulations biasing the LDA coordinate. Here, we will test three different hill heights (0.5, 0.9, 1.3 kcal/mol) to check the convergence. In the included simulations, we also use walls at +20 and −30 (commands omitted below for brevity) to prevent the system going far outside physical ranges of the LD variable seen in Fig. 5.

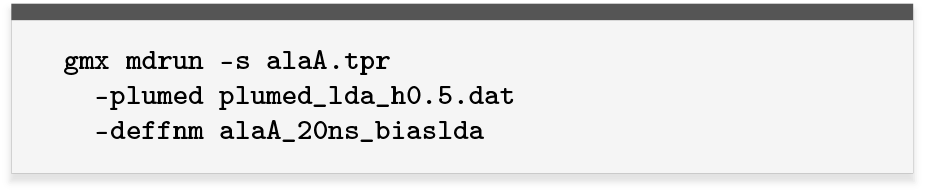

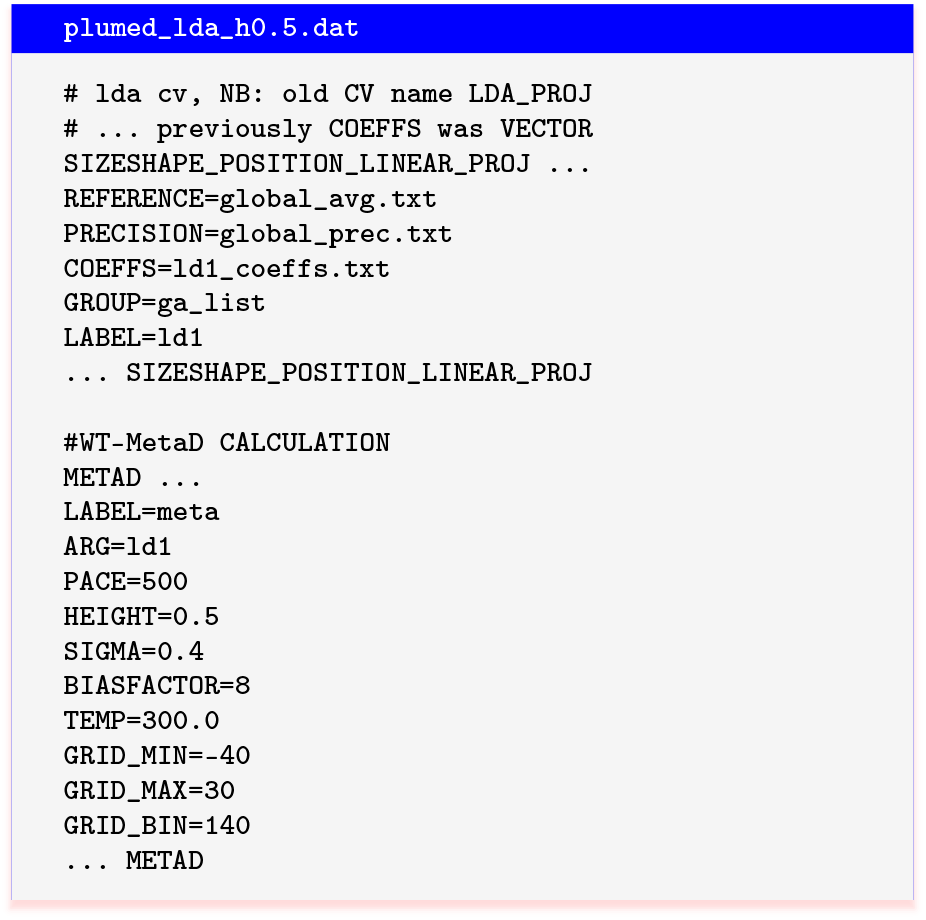

We first confirm that the LD coordinate is going back and forth between the values we expect for our two different states (Fig. 6, top). We also check that the slow *ϕ* degree of freedom for ADP is being sampled (Fig. 6, bottom).

**FIG. 6.**
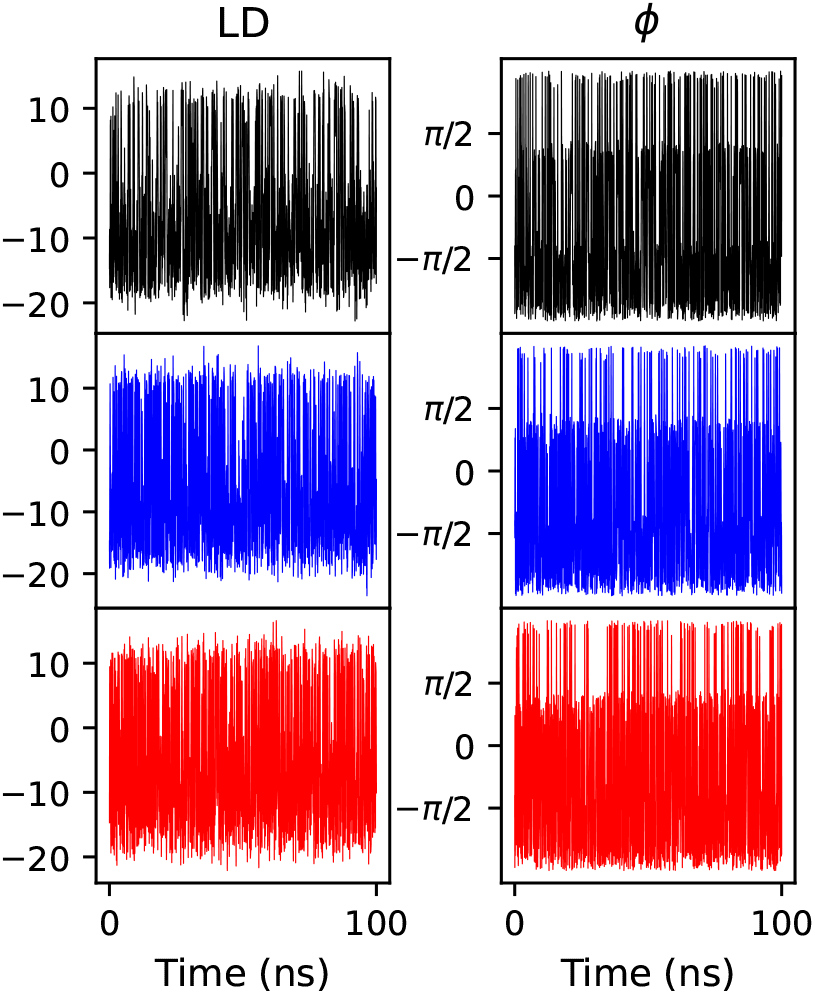
LDA coordinate value and *ϕ* coordinate versus time in three WT-MetaD runs. The three runs differ in MetaD height *h* =0.5 (black), 1.3 (blue), or 1.7 (red) kcal/mol.

In a real/larger biomolecular problem, we have found is important to check that the true end states are being accessed and not only that those LD values are observed, since the LD transformation need not be bijective. We also note that the quality of sampling can certainly depend on the amount of available input data, which defines the cluster means and covariances as well as the global alignment. This is addressed both in our original biasing study^7^ as well as our work on iteratively improving these quantities through further biased sampling (next section).^8^

We can compute the FES from these runs projected in Ramachandran space (Fig. 7 left). We note that sampling of the FES appears visually correct. We can make a quantitative comparison by comparing these FES bin-by-bin with the reference FES in Fig. 1, as shown in the right column of Fig. 7.

**FIG. 7.**
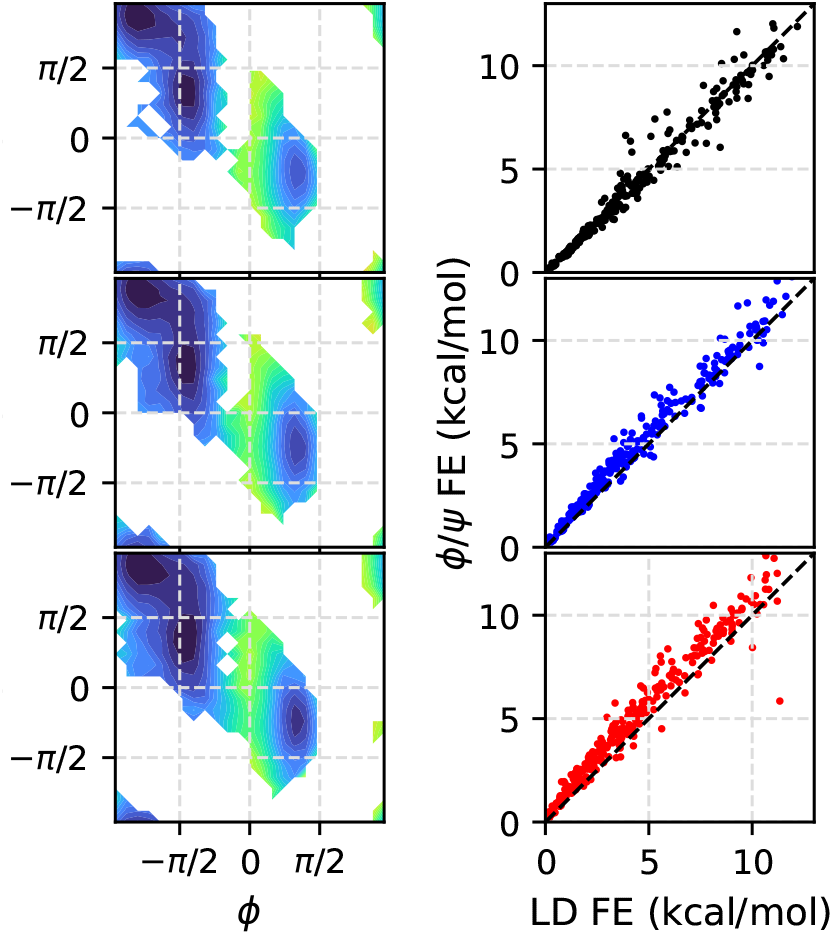
FES in Ramachandran space from MetaD performed on LDA coordinate with three *h* values: 0.5 (top/black), 1.3 (middle/blue) and 1.7 (bottom/red) respectively as shown in Fig. 6. The color scale is the same as Fig. 1, with energies from 0 to 11 kcal/mol. Right is a bin-by-bin comparison of the left column with the values in Fig. 1.

Finally, we note that we can also check whether the MetaD is converging. We do this by computing the FES at intermediate times in the MetaD run along the biased coordinate (Fig. 8). We can also compute the FES along this coordinate for the reference *ϕ* /*ψ* MetaD simulation as a comparison. In this case, the FES converges rapidly to the reference, although it appears to be less accurate for faster biasing.

**FIG. 8.**
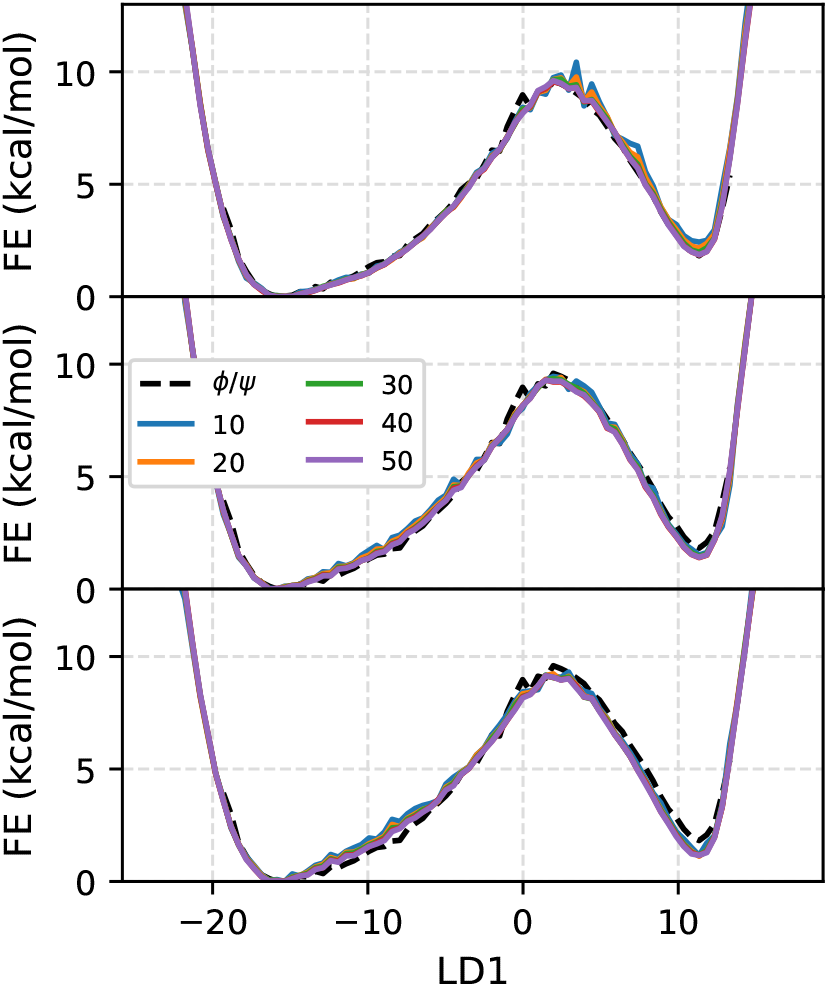
FES computed along LDA coordinate at different intermediate times (10ns to 50ns with an interval of 10ns), obtained from MetaD simulations performed at three *h* values: 0.5 (top), 1.3 (middle) and 1.7 (bottom) respectively. The FES convereges quickly to the reference dashed line.

**FIG. 9.**
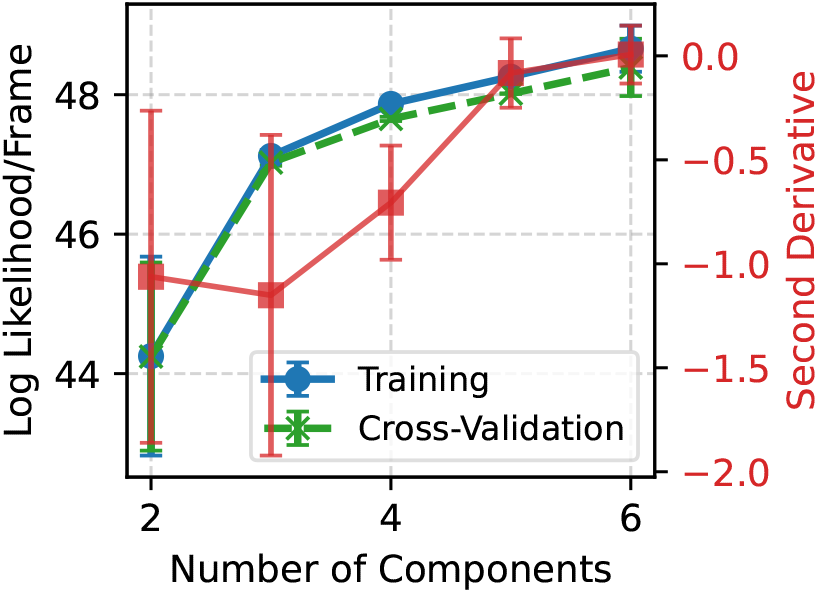
Same as in Fig. 3, except now bias weights for each frame have been included in the fit.

**FIG. 10.**
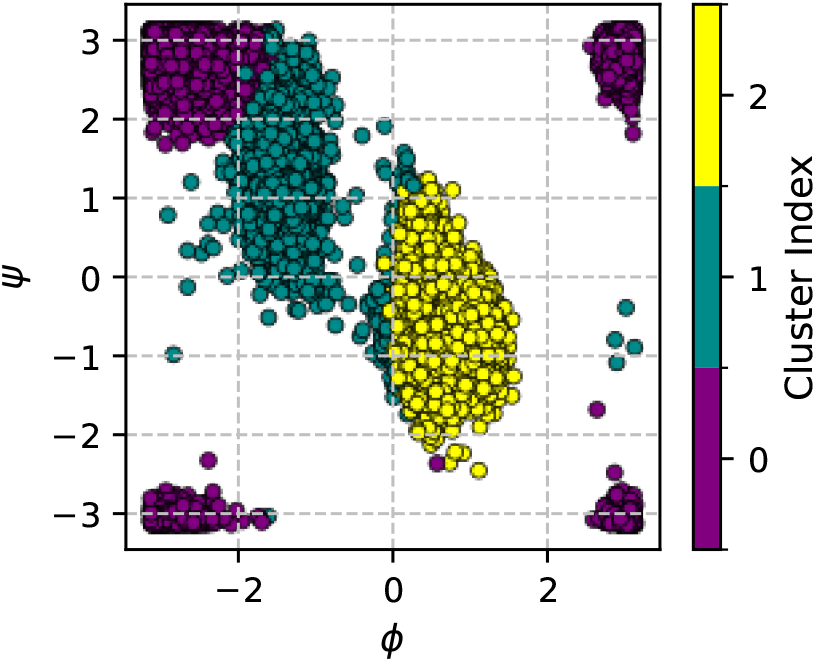
Ramachandran plot with points colored by *K* = 3 clustering on the data generated by MetaD on an LDA coordinate.

### F. Clustering on biased data

We now will attempt to cluster output of our WT-MetaD simulation, where we biased the LDA coordinate using a hill height of *h* = 0.5 kcal/mol. This just requires a small change from our procedure in Sec. IV C. It should be noted here, that we are providing the weights for training data in ShpaeGMM fit call in addition to the coordinate from the WT-MetaD trajectory. The weights are computed using the Eq. 11. From the minimum in the second derivative, we take *K* = 3 to proceed to the next step.

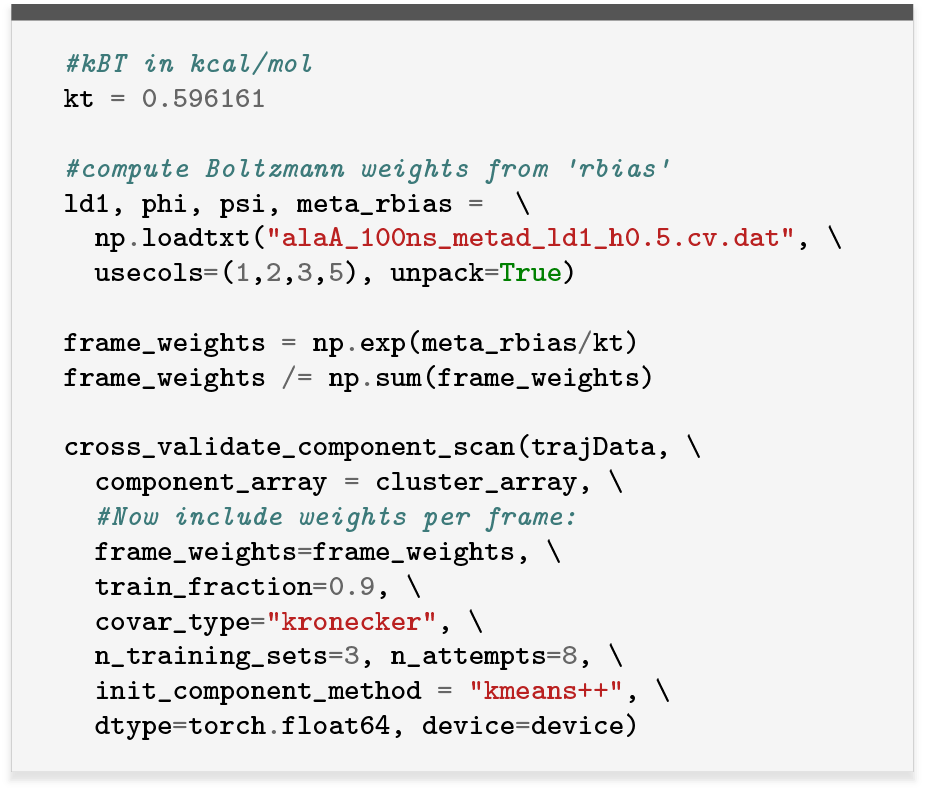

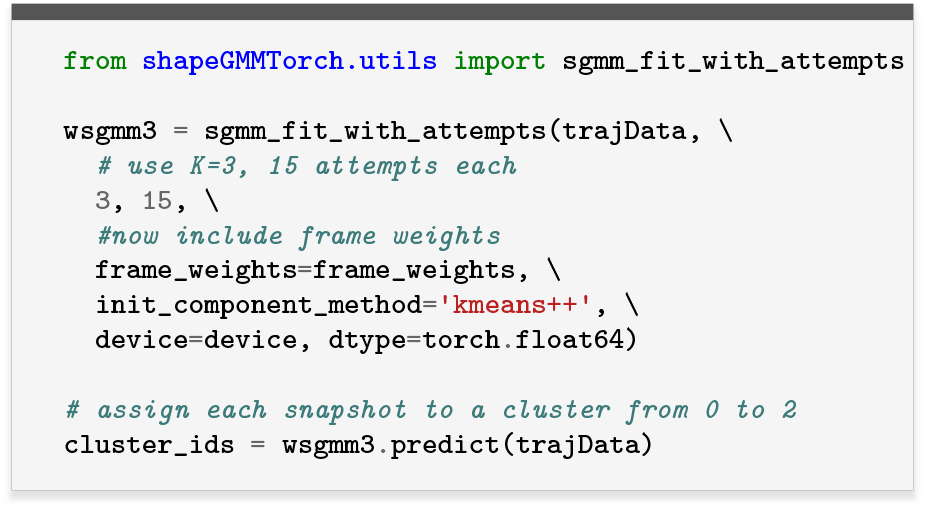

Finally, if we want, we can fit a new LDA coordinate with these weighted data as done in Ref. 8. To do this, we compute the Bhattacharya distance between the new and old cluster definitions, and take the new clusters that are at minimum distance from the old ones. The results are shown in Fig. 11.

**FIG. 11.**
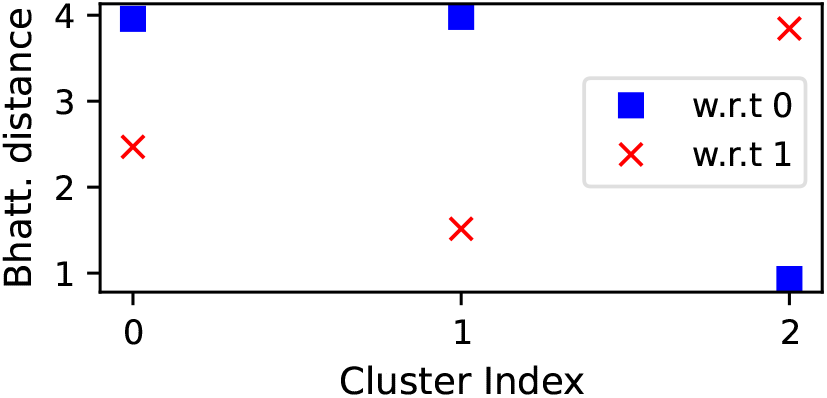
Bhattacharya distance between our new and old cluster definitions. The new cluster 2 corresponds to the old cluster 0, and the new cluster 1 corresponds to the old cluster 1.

**FIG. 12.**
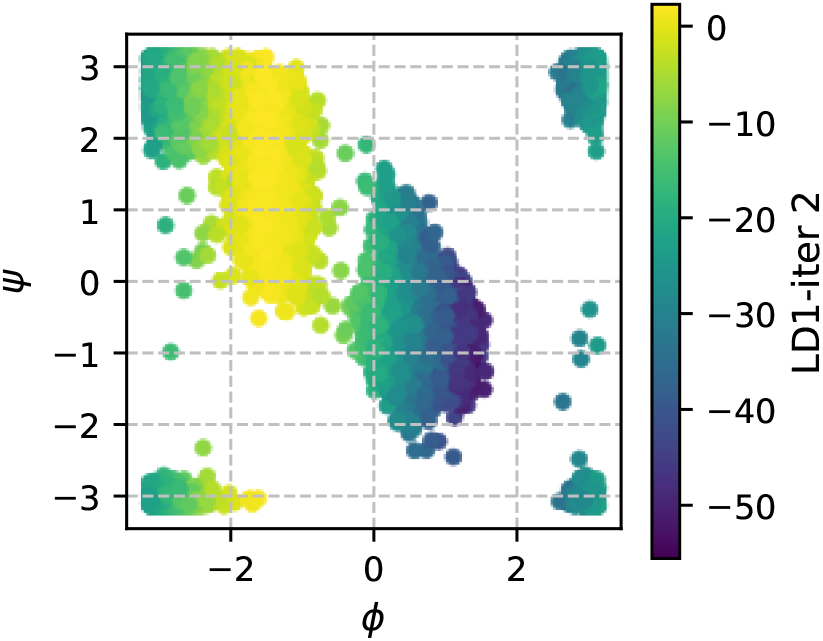
Ramachandran plot with points colored the value of our new LDA coordinate.

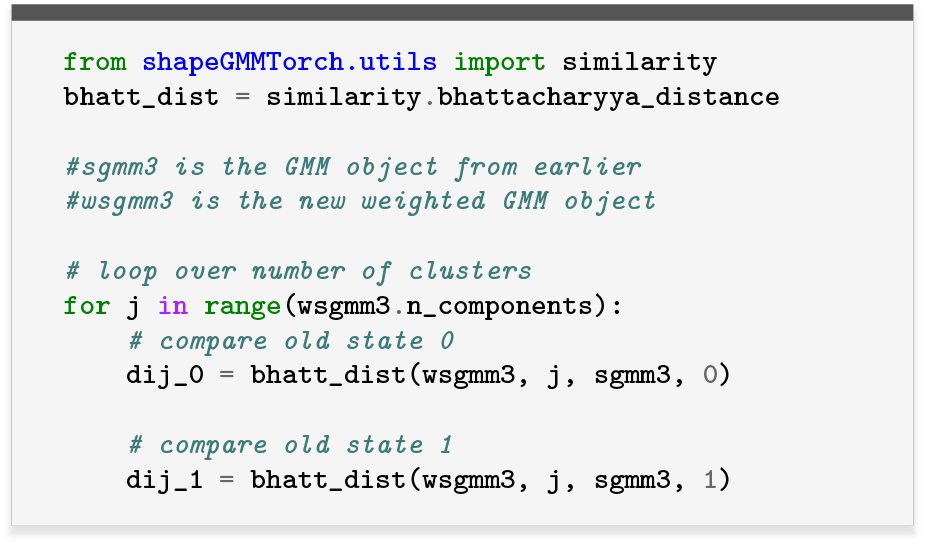

We now repeat our global alignment procedure from earlier, replacing uniform frame weights with the values from MetaD. After that, we perform our LDA fit as before, again using frame weights.

If we project the value of the new LDA coefficient on these points, we see that the coordinate does separate the basins in a continuous way as desired.

### G. Generating samples from the model and estimating thermodynamics quantities

Our GMM objects are probabilistic models. We can therefore generate samples and use these samples to estimate thermodynamic quantities. As one example, we show how we can estimate the FES, which can in this case be compared to our reference surface from MetaD on Ramachandran angles. In Fig. 13, the generated points are shown on the left, with the estimated equilibrium FES on the right, and the reference surface shown as contours.

**FIG. 13.**
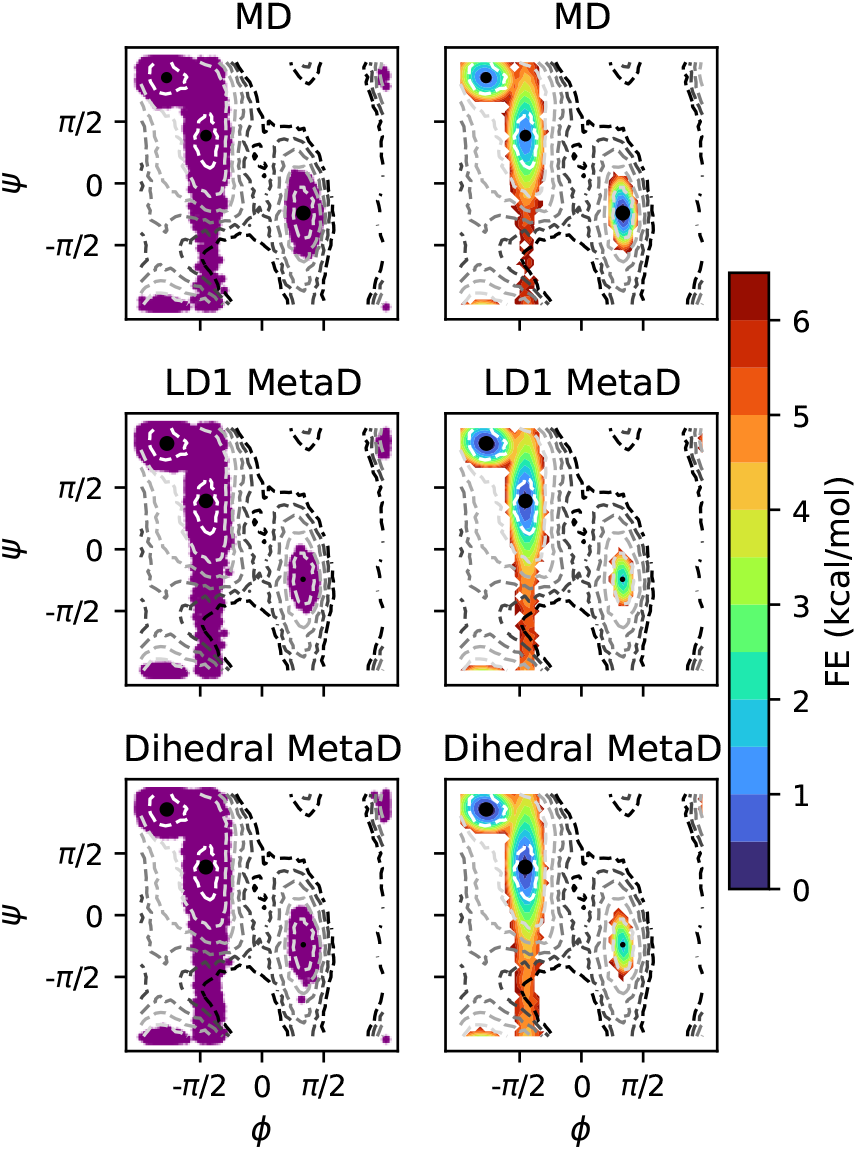
Results from generating samples using three different ShapeGMM models, fit on equilibrium data, MetaD data on LDA, and MetaD data on Ramachandran angles (reference data). Left shows the generated points, and right the FES in Ramachandran space. The dashed contours show the reference FES.

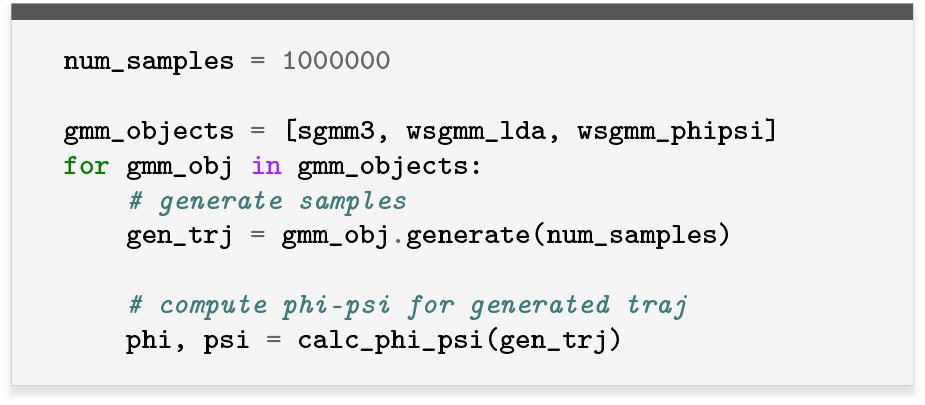

Our ability to generate points allows us to estimate thermodynamic quantities, including the FES shown in Fig. 13. As another example, we can compute the (relative) configurational entropy as described in Sec. III C 1. To illustrate this point visually, we train a three-state model on each of the short MD trajectories shown in Fig. 2. Intuitively, we expect the entropy of the ensemble sampled by these short MD trajectories to be much smaller than that of the full ensemble calculated using MetaD.

Our simple procedure for computing the configurational entropy is given below, with results shown in Fig. 14. As expected, models trained on data from just one basin generate samples only in that region, and entropies computed from these models are smaller than the reference. The difference in entropy is more negative for the smaller basin.

**FIG. 14.**
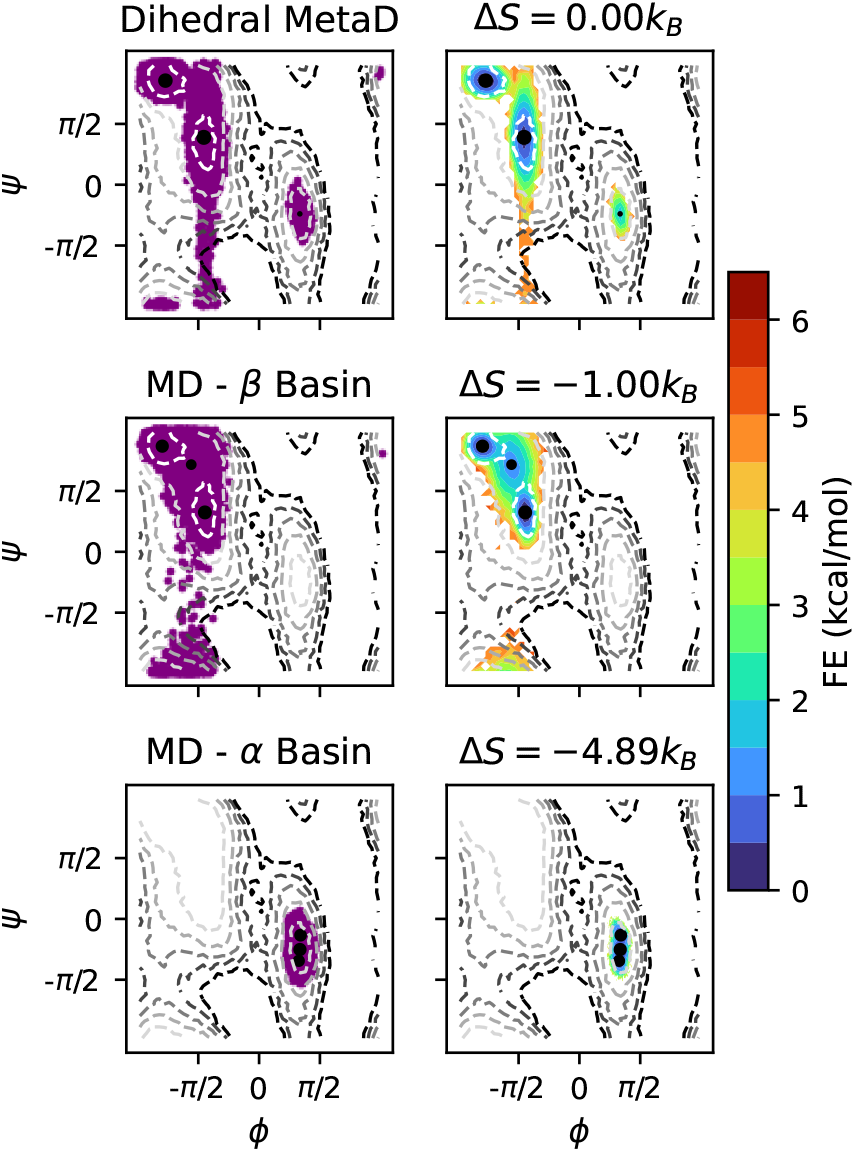
Results from generating samples using three-state ShapeGMM models fit on MetaD data, and on equilibrium MD started in two different basins. Titles on right report the difference in entropy between the models and the reference model (top row). Contours and colorbars same as Fig. 13.

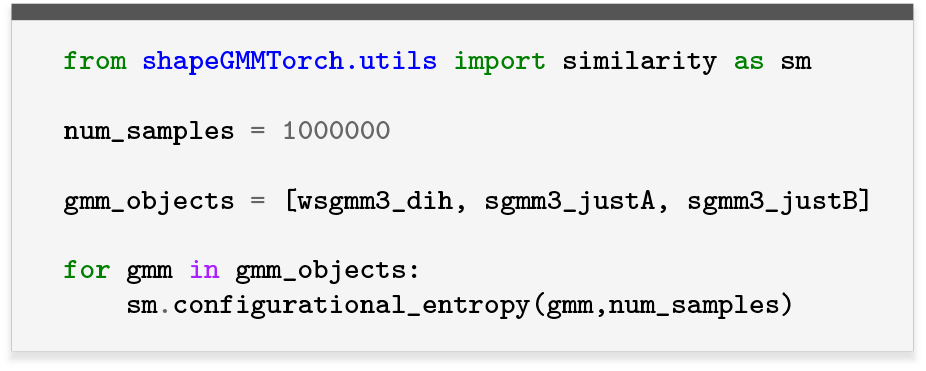

## V. CONCLUSIONS

In this tutorial, we have outlined a complete workflow for applying the ShapeGMM for clustering, characterizing, and sampling conformational ensembles of biomolecules. By working in size-and-shape space, ShapeGMM provides a principled way to use Cartesian atomic positions as features while remaining invariant to rotation and translation. We demonstrated how to fit probabilistic models to unbiased trajectories, extract thermodynamic information, and define reaction coordinates that can be directly employed in WT-MetaD simulations. We further showed how biased simulations can be incorporated back into ShapeGMM fits through appropriate reweighting, enabling the generation of refined equilibrium models even when enhanced sampling is required to access relevant states.

Although illustrated on the alanine dipeptide system, the procedures presented here are readily transferable to larger and more complex systems where unbiased sampling is often infeasible. For system sizes above ∼ 200 atoms, users must be careful about the curse of dimensionality as well as the increased computational cost of the fitting (see Sec. VII for additional insight). For these systems, it is typical to select a subset of atoms, e.g. CA atoms, that should adequately describe the coarse-grained motion. We have successfully applied ShapeGMM with Kronecker covariance to SARS-CoV-2 nsp13 using ∼440 CA atoms from the catalytic domains of the protein^9^. For system sizes much larger than this, further approximations may need to be made such as additional coarse-graining of the atom selection, selecting a subset of the protein such as we did for WNV NS3,^10^ or using a uniform covariance model which has significantly fewer parameters than the Kronecker covariance model. Regardless, the accompanying software environment, worked examples, and containerized installations along with Jupyter notebooks make it straightforward for researchers to reproduce our results and adapt the workflow to their own problems. Beyond clustering and free energy estimation, ShapeGMM has already found applications in ensemble reweighting, kinetics,^35^ and the development of improved collective variables through iteration on biased data,^8^ highlighting its versatility as a general framework for analyzing molecular simulations.

We anticipate that this methodology will aid researchers in both extracting mechanistic insight from large-scale molecular simulations and in designing enhanced sampling protocols guided by statistically principled models. While others have already used ShapeGMM package,^36,37^ we hope that this tutorial will help lower the barrier for the broader use of ShapeGMM in our scientific community, providing a statistically rigorous and practical approach to quantifying and sampling conformational ensembles in biomolecular systems.

## VI. APPENDIX A: USING PROVIDED CONTAINER FILES

We have generated and provide a Docker/Singularity container with all functionality required to perform the steps of this tutorial. We have also included a Dockerfile that lists every step used for fully installing the necessary software.

To obtain the appropriate container, one can run

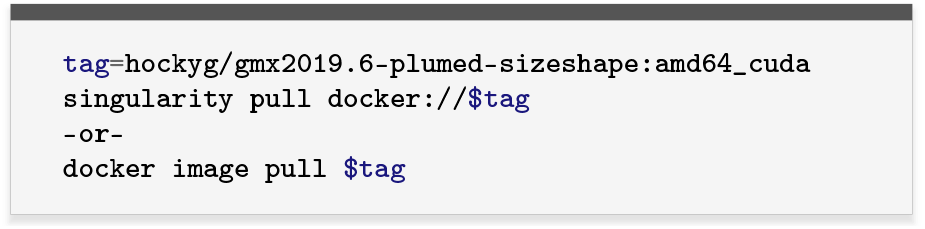

One can then use Docker or singularity to run commands:

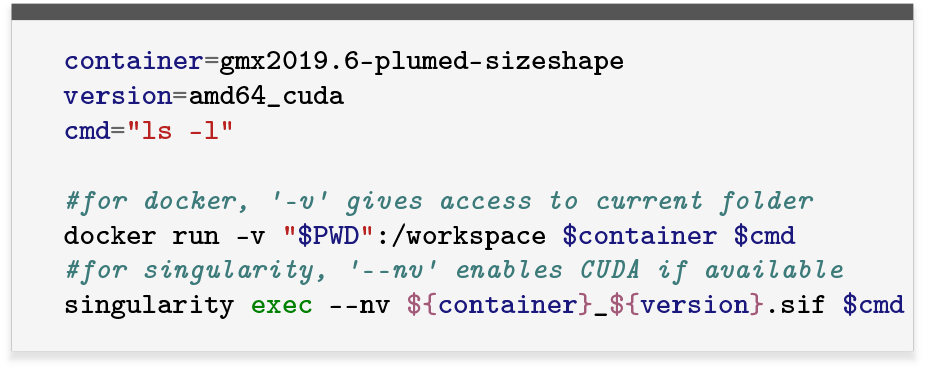

Alternatively, one can use the tag `hockyg/gmx2024-plumed:amd64_cuda` to obtain a container with GROMACS 2024.3 and Plumed 2.10b. This container has all available PLUMED modules installed, including `sizeshape`.

## VII. APPENDIX B: COMPUTATIONAL COST OF SHAPEGMM

The `shapeGMMTorch` code uses `PyTorch` linear algebra routines and thus performs well on both the CPU and GPU. Here we assess the performance of the GPU code on a single NVIDIA RTX 4090 GPU. As a test system, we use the 305 microsecond trajectory of the fast-folding mutant of HP35 from DE Shaw.^38^ We have previously used this system as a benchmark for clustering^4^ as well as reaction coordinate determination.^7,8^ The computational cost of determining optimal ShapeGMM parameters depends on five main factors: (1) the covariance type, (2) the number of atoms/particles, (3) the number of components, (4) the training set size, and (5) initial conditions. Here we use only the `kronecker` covariance type which is more computationally demanding than the `uniform` covariance. Additionally, we restrict the training set size to 100k frames for simplicity.

We find that ShapeGMM scales linearly over components and number of atoms on a single GPU for the range of number of atoms (36 to 455) and components (1 to 6) chosen. The ShapeGMM optimization time for every combination of number of atoms and number of components was averaged over 20 randomized initializations. The linear behavior over number of components is not shown but the slope of these fits yields the minutes per component optimization. This value for each choice of the number of atoms is plotted in Fig. 15. The linear behavior of this plot suggests that the parallelization of the code on the GPU is efficient over this atom range because a naive calculation of the covariance would scale quadratically. For 250 atoms, for example, it takes approximately 0.25 minutes per component optimization. In other words, it would take one minute to optimize a 4 component model with 250 atoms. We note, however, that we typically optimize 5-10 models (attempts) per hyperparameter set and choose the highest log-likelihood model to attempt to avoid local maxima. Regardless, this scaling suggests it is feasible to fit ShapeGMM models for system sizes under 500 atoms in under an hour.

**FIG. 15.**
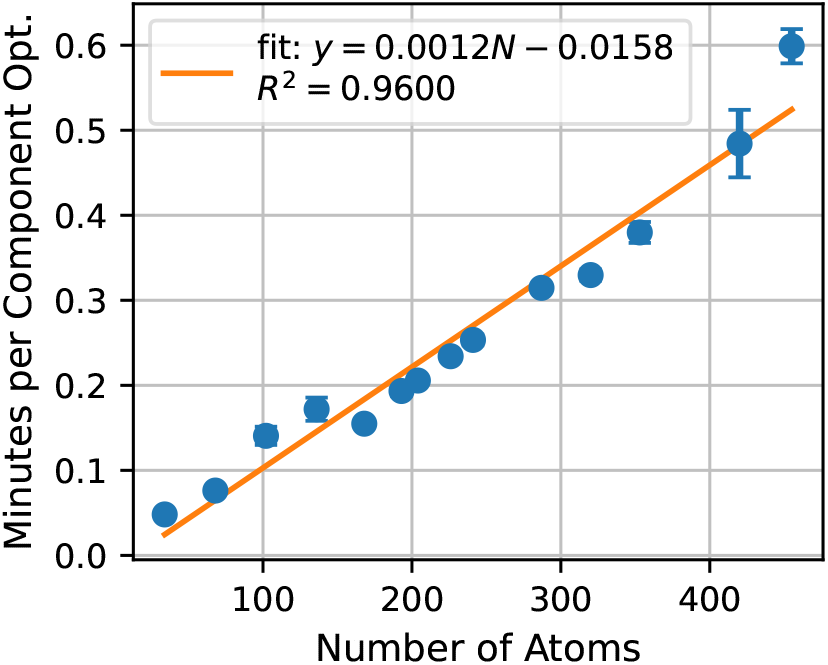
Computational performance of ShapeGMM as a function of number of atoms for 100k training frames from a trajectory of HP35. The computational performance is assessed as the minutes per component optimization (slope of the minutes to optimize as a function of number of components). This was assessed on a single RTX 4090 GPU.

## ACKNOWLEDGMENTS

SS and GMH were supported by the National Institutes of Health through the award R35GM138312. SS was also partially supported by a graduate fellowship from the Simons Center for Computational Physical Chemistry (SCCPC) at NYU (SF Grant No. 839534). MM acknowledges funding from National Institute of Allergy and Infectious Diseases of the National Institutes of Health under award number R01AI166050 and the National Science Foundation under award 2238706. MM would like to acknowledge Peter T. Lake, who implemented the frame-weighted LDA method as a Python package.^14^ This work was supported in part through the NYU IT High Performance Computing resources, services, and staff expertise, and simulations were partially executed on resources supported by the SCCPC at NYU.

## DATA AVAILABILITY STATEMENT

All data and code for this tutorial are available from a GitHub repository for this article: https://github.com/hocky-research-group/ShapeGMM_tutorial_2025.

